# Finding Fluorescence: Utilizing community science to document novel biofluorescence occurrences and encourage community engagement in science

**DOI:** 10.1101/2024.04.24.590905

**Authors:** Hannah Burke, Lauren Serrano, Emily Lemmon, Courtney Whitcher

**Author notes:** Corresponding author (HB).

## Abstract

Fluorescence, a form of photoluminescence, is the emission of light at a longer wavelength by a substance when exposed to shorter-wavelength energy. Biofluorescence, which can be observed in living organisms, involves the absorption of light at one wavelength and re-emission at a longer wavelength due to fluorophores in specialized cells or structures. While initially studied in marine organisms, attention has shifted to exploring biofluorescence in terrestrial organisms, revealing roles in reproduction, camouflage, communication, and prey attraction across phyla. Community science databases engage the public in data collection, fostering scientific discovery and strengthening the science-society connection. Such databases have become valuable tools and have aided scientists in understanding the natural history of many different traits in organisms. This paper introduces *Finding Fluorescence*, the first biofluorescence-based community science website established in 2020 to gather public observations of biofluorescent organisms. The study presents at least 15 novel biofluorescence accounts spanning five phyla, 15 families, and 15 species. The observations collected from *Finding Fluorescence* contribute to our understanding of fluorescence in organisms and provide insight into possible ecological functions. We emphasize the importance of community engagement in scientific exploration and encourage future studies to incorporate such aspects into their research.

## Introduction

Fluorescence is a form of photoluminescence which occurs when a substance emits light at a longer wavelength after exposure to an energy of shorter wavelength [1]. Biofluorescence, more specifically, is the phenomenon in which a living organism absorbs light at one wavelength and re-emits it at a longer wavelength [2]. On a chemical level, biofluorescence involves fluorophores inside the specialized cells or structures of organisms [3]. Exposure to short-wavelength light (e.g. UV light) causes these fluorophores to absorb the energy from the photons and enter a higher energy state [3]. Excess energy as visible light (i.e. the fluorescence observable to the eye) is emitted when the photon returns to a lower energy state [3]. The phenomenon of fluorescence was initially studied in marine organisms, as catalyzed by the discovery of the green fluorescent protein (GFP) in the cnidarian *Aequorea victoria*, now commonly used to fluorescently mark genes in organisms [4]. However, attention has recently turned to examining fluorescence presence as well as possible ecological significance in terrestrial organisms. Biofluorescence is found across all major phyla of terrestrial organisms and has been found to play a role in key biological processes such as reproduction, camouflage, communication, and prey attraction [5]. Documenting new fluorescence presences is one key step toward beginning to better understand the ecological role fluorescence may have across taxa. As a quick reference for understanding biofluorescence throughout the paper, a simple vocabulary aid has been included (Table 1).

**Table 1.** Fluorescence vocabulary.

| Term | Definition |
| --- | --- |
| Fluorescence | Fluorescence is an optical occurrence in which an object emits light at a longer wavelength when exposed to light at a shorter wavelength [1]. |
| Biofluorescence | Biofluorescence is the reemission of longer, lower-energy wavelengths after absorbing photons in biological tissues [6].<br>Biofluorescence and fluorescence will be used interchangeably throughout this paper, as we are only examining fluorescence in living organisms. |
| Excitation | Excitation refers to the wavelength of light used to induce fluorescence in an organism [3]. In the paper, it will commonly be used to describe the wavelength of light that was shone on each community science submission. (E.g. An organism fluoresced when *excited* by a wavelength of 400 nm). |
| Emission | Emission refers to the wavelength of light that is fluorescing from an organism when exposed to its specific excitation range [5]. In this paper, emission will be characterized simply by the color emitted from the organism during excitation (E.g., the mushroom showed a green, fluorescent *emission* when excited by 400 nm light). |
| Fluorophores | Fluorophores refer to chemical compounds inside specialized cells or tissues in organisms which are known to fluorescence when excited [3]. Fluorophores are the chemical explanation for fluorescence in organisms. |

Common terminology related to fluorescence that is seen throughout the paper for quick referencing to terms.

With so many organisms having never been examined for fluorescence, there is much ground to cover, and community science databases have the potential to provide broad discovery and insight. Databases that function as community science projects are excellent methods to facilitate large-scale data collection efforts efficiently and affordably. For example, one study examined the migration of 15 waterfowl bird species by building models from 43 years of community science data; these models revealed that precipitation and minimum temperature significantly influenced migration phenology for most species being considered [7]. Community science databases allow submissions (e.g., photographs, written observations, measurements, etc.) from the public which offer an expansive pool of information for researchers across many disciplines. As examples of the raw potential these submissions hold, prior studies have employed submissions such as these to inform land management decisions [8]; document impacts of climate change [9]; and address questions related to species distributions [10], species interactions [11], and species health [12]. Needless to say, community science has the potential to reach a broad range of science and holds much promise for future studies as well as conservation efforts.

These databases allow the public to participate in data collection and analysis, thereby accelerating scientific discoveries [13]; fostering public engagement [14]; empowering individuals to actively contribute to scientific endeavors [15]; and promoting a deeper connection between science and society [16]. Interpreting these databases commonly involves validation processes to ensure data accuracy and reliability [14]. Researchers employ various statistical methods and tools to analyze the information, uncovering patterns, trends, and insights from the community submissions to satisfy their respective goals [14]. Clear guidelines and instructions are additionally provided within community science databases to avoid as many submission errors as possible and to ensure the uploads are consistent with the project’s desired formatting [14].

*Finding Fluorescence* is a public education and community science website designed in 2020 to encourage individuals to submit their own observations of organisms they find that fluoresce [17]. *Finding Fluorescence* aims to encourage scientific exploration and public interest in organisms present in the everyday environment while enabling a collaborative effort to collect vast amounts of novel scientific data on the presence of biofluorescence across taxa. We sought to aid future studies examining fluorescence in organisms by presenting novel community science findings of biofluorescence across various taxa found via submissions sent to *Finding Fluorescence*.

Here we present at least 15 new accounts of biofluorescence, spanning 15 species, 15 families, and five phyla, all of which were found by community scientists. A new account of biofluorescence was documented if a new variation in fluorescence color, fluorescence pattern, or excitation wavelength inducing fluorescence was observed in each organism. After analyzing the submissions to our community science database, we propose additional improvements to the site which may help facilitate better community science data collection in the future.

## Methods

### Project design

Our project aimed to assess the effectiveness of the *Finding Fluorescence* community science resource as well as document new findings of fluorescence. To do so, we assessed the observations uploaded to the community science database. For each observation we (1) verified the data, (2) determined if the observation was novel, and (3) documented future improvements and directions for submission criteria. The data utilized in this study was retrieved from the *Finding Fluorescence* community science database. Each uploaded *Finding Fluorescence* observation contains information under the following headings: *date; time; location; type of organism; common and/or scientific name; is there fluorescence; description of fluorescence (location, pattern, color, etc.); wavelength of light used (or model of light if wavelength unknown); any other information (weather, habitat, behavior of organism, etc.); a photo of the fluorescence; and a photo of the organism in natural (or white) light,* as defined by the data upload form implemented via Epicollect5 [18]. On 3 December 2023, we downloaded all observations submitted since the launch of the site in 2020 and verified the data for accuracy, with the aim to identify to at least the family level. After data verification, we conducted an extensive literature review to determine if the observations uploaded to the site were novel or if prior accounts of biofluorescence were previously recorded in that taxa.

### Organism verification

We verified the identification of each submission by employing Florida State University’s Library Database [19], Google Scholar [20], and various field guides and community science resources found via Google’s search engine [21]. In the Florida State University (FSU) Library Database, we applied advanced search and entered the submitted scientific/common name and selected AND to include “field guide” in the search. If no scientific or common name was submitted, we utilized the description of the organism along with its location and the uploaded photographs to aid in identification. We also verified organism identity through Google and Google Scholar with the previous procedure if FSU’s Database did not provide any results. We then compared the description and/or photograph of the organism found via our literature search to the description/photograph submitted to *Finding Fluorescence* to confirm its scientific and/or common name. To gain further confidence in our proposed identifications for various taxa, we collaborated with biologists within Florida State University (FSU) to reinforce our identification proposals. For observations where a family-level verification was not possible or confident, we chose to still include under an *Unidentified Taxa* subheading in the paper.

### Prior relevant biofluorescence literature review

After identification, we searched for literature documenting prior fluorescence of that organism or the fluorescence of a closely related species for each submission (i.e., for an organism identified down to the species level, another organism within the same genus would be a closely related species. If no organism in the same genus was found to fluoresce, the search would be extended to an organism in the same family). As stated prior, we used FSU’s Library Database, Google Scholar, and Google as search engines for prior recorded biofluorescence accounts. For FSU’s database, we included the verified species name (if possible) along with the terms “biofluorescence” OR “fluorescence” to allow a broad search of publications that discuss their biofluorescence. We repeated this procedure for both Google and Google Scholar. If no results were returned, we then proceeded to generalize the search terms to genus and then to family to find whether a closely related organism had prior fluorescence documentation. For any record of fluorescence description found, we recorded the excitation wavelength, organism, and description of fluorescence, as available. It must be noted that we were only interested in examining prior natural fluorescence documentations in taxa. We were not including studies that were using fluorescent labeling or manipulating fluorescence in the organism. The fluorescence we are interested in for this study and for prior relevance is fluorescence that can be seen from the organism itself after induced excitation (such as with a UV flashlight).

### Data availability and sharing

All biofluorescence observations can be found on the public community science database Finding Fluorescence at https://findingfluorescence.wixsite.com/home/view-map.

## Results

### Global impact of Finding Fluorescence

The breadth of website engagement (i.e. any type of access to the site) for *Finding Fluorescence* reached 64 different countries, 48 different states within the United States, and six out of the seven continents in the world (Fig. 1). Submissions from community members came from three different countries around the world, including the United States of America, the Netherlands, and Belize (Fig. 2). We received a total of 36 submissions, with 32 submissions from the United States of America, three from the Netherlands, and one from Belize. Within the United States, we received submissions from Florida, Michigan, New Jersey, Pennsylvania, and Illinois. Observations collected from ecologically diverse sites within the United States and around the world allowed for an array of organisms to be assessed.

**Fig 1.**
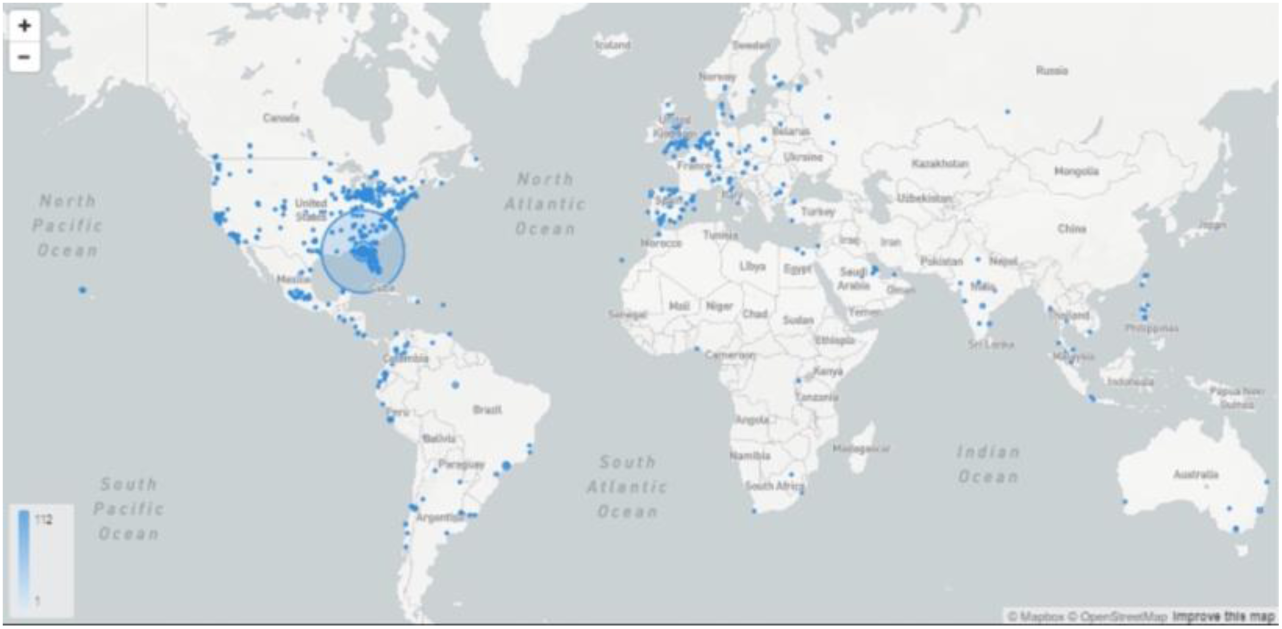
Global *Finding Fluorescence* engagement. The current reach of the *Finding Fluorescence* resource. Blue points indicate interaction with at least one of the materials provided on https://findingfluorescence.wixsite.com/home.

**Fig 2.**
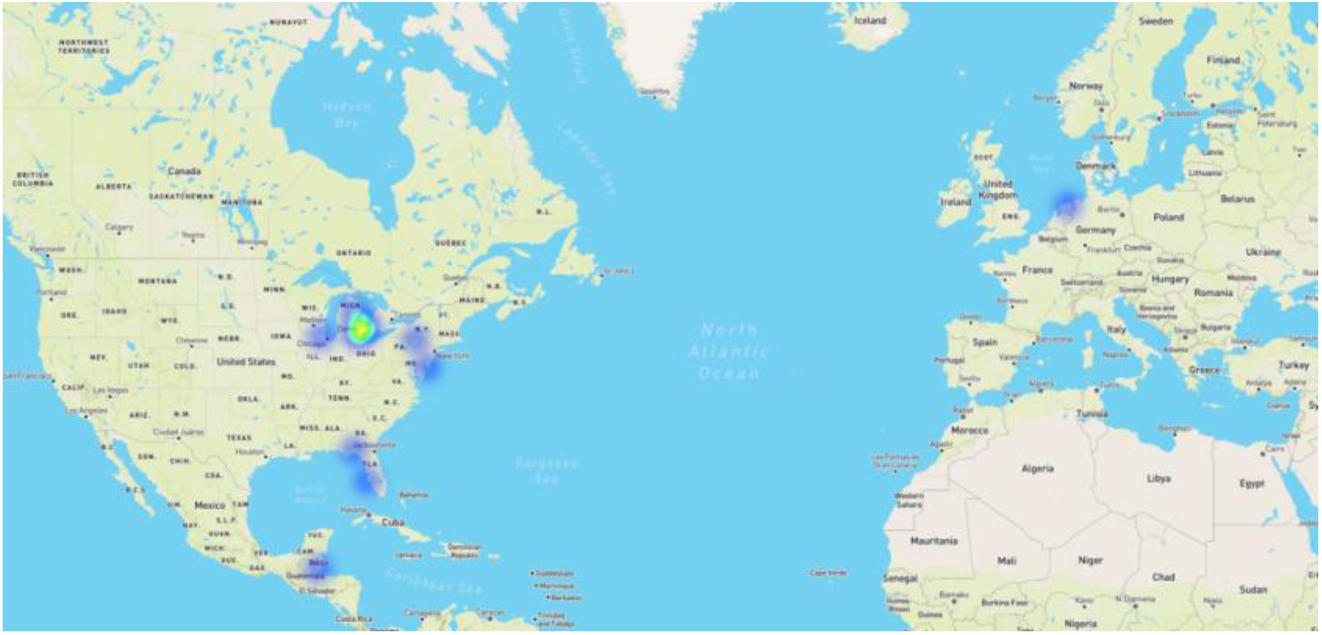
Submissions to *Finding Fluorescence* by location. The current locations of where submission to *Finding Fluorescence* originated from. Areas of yellow indicates concentrated submissions from a particular area, while blue indicates at least one submission from a given location.

### Documenting new discoveries of biofluorescence

All descriptions below are documents of fluorescent observations uploaded to the *Finding Fluorescence* community science database (Table 2). These observations have been categorized under their respective phylum. Any observation that is not entirely new (i.e., there has been a prior fluorescence documentation of a related organism up to the family level) is also included (Table 3) and will be addressed later in the paper. The wavelengths included in our results were self-reported by the community members who submitted the fluorescence documentation. If the exact fluorescence excitation was unknown — either because no specific excitation flashlight model was included, or no wavelength was stated — we simply listed the possible range of light that could have been used given the supplied information. As well, some observations list potential fluorescence documentations, as we were unable to verify nor deny fluorescence observations given the nature of some of the submitted photographs.

**Table 2.**
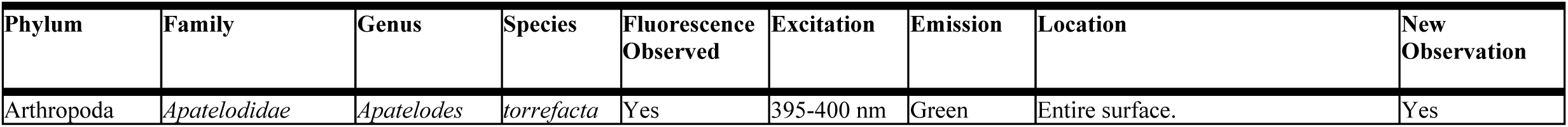
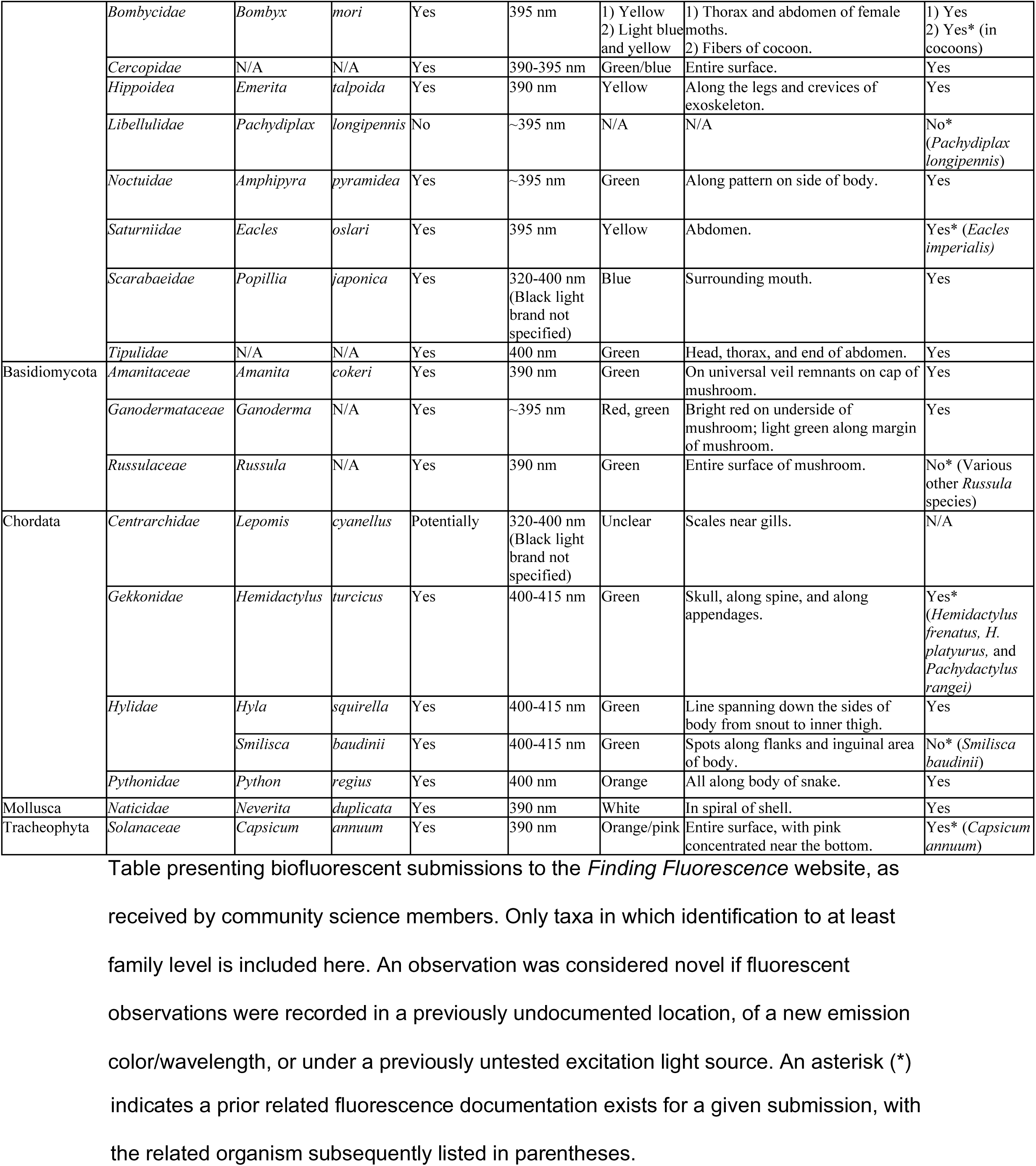
Verified *Finding Fluorescence* submissions.

**Table 3.** Prior relevant fluorescence documentations.

| Phylum | Family | Genus | Species | Fluorescence Observed | Excitation | Emission | Location | Source |
| --- | --- | --- | --- | --- | --- | --- | --- | --- |
| Arthropoda | *Bombycidae* | *Bombyx* | *mori* | Yes | 1) 360 nm<br>2) 365 nm | 1) Yellow-white and blue-purple<br>2) Yellow and blue | 1) Cocoon fibers.<br>2) Cocoon fibers. | 1) [22]<br>2) [23] |
|  | *Libellulidae* | *Pachydiplax* | *longipennis* | Yes | 310-340 nm | Blue | Thorax, along wing hinges. | [24] |
|  | *Saturniidae* | *Eacles* | *imperialis* | Yes | 320-400 nm | Orange and brown-lavender | Not specified. | [25] |
| Basidiomycota | *Russulaceae* | *Russula* | Various *Russula* spp. | Yes | 320-400 nm (Black light brand not specified) | N/A | N/A | [26] |
| Chordata | *Gekkonidae* | *Hemidactylus* | *frenatus* | Yes | 395-400 nm | Green | Skull, along spine, and along appendages. | [27] |
|  |  |  | *platyurus* | Yes | 395 nm | Blue | Along the lower jaw bone. | [28] |
|  |  | *Pachydactylus* | *rangei* | Yes | 365 nm | Green | Skin around eyes and flanks. | [29] |
|  | *Hylidae* | *Smilisca* | *baudinii* | Yes | 400-415 nm | Green | On arms and inguinal area. | [30] |
| Tracheophyta | *Solanaceae* | *Capsicum* | *annuum* | Yes | 1) 365 nm | 1) Blue | 1) Uniformly distributed on surface | 1) [31] |
|  |  |  |  |  | 2) Maximum 440 nm | 2) Blue, low-green, low chlorophyll | 2) Not specified | 2) [32] |

Table displaying prior fluorescence documentation found in related organisms. We
considered an organism to be relevant if it was found to have fluoresced and was within the same family of an organism submitted to *Finding Fluorescence*.

### Phylum Arthropoda

An observation revealed that the spotted Apatelodes moth caterpillar (*Apatelodes torrefacta*) displayed whole body biofluorescence, which appeared as a bright green color under an excitation light of 395-400 nm. In white light, the spotted Apatelodes displayed a yellow color on its thorax and abdomen. The domestic silk moth (*Bombyx mori*) displayed yellow fluorescence on the abdomen of female specimens under an excitation wavelength of 395 nm. Under white light, the abdomen of the moth was white. Fibers from the cocoon of *B. mori* displayed yellow and blue fluorescence separately (i.e. one cocoon displayed blue fluorescence only and another displayed yellow fluorescence only) on cocoons at the same excitation wavelength of 395 nm*. Under white light, the cocoon fibers were white. Frog hoppers, also known as spittle bugs (family *Cercopidae*), fluoresced when exposed to 390-395 nm wavelengths. The fluorescence appears as a blue-green emission on the arthropod’s entire surface, while in white light the frog hoppers appeared as a white/creamy color. The Atlantic mole crab (*Emerita talpoida*) revealed a yellow fluorescence on the legs and crevices of the exoskeleton under an excitation wavelength of 390 nm. Under white light, the mole crab displayed a light brown and white pattern. The blue dasher dragonfly (*Pachydiplax longipennis*) did not show fluorescence under excitation wavelengths of 395 nm*. The copper underwhig moth caterpillar (*Amphipyra pyramidea*) exhibited green fluorescence along its patterns on the lateral sides of its body under an excitation light of 395 nm. In white light, the underwhig caterpillar was a uniform light green color with yellow patterns on its sides. The Oslar’s eacles moth (*Eacles oslari*) fluoresced neon yellow on its abdomen under an excitation wavelength of 395 nm. Under white light, the abdomen of the moth was a mix of yellow and brown markings. The Japanese beetle (*Popillia japonica)* displayed blue fluorescence near its mouth with an excitation wavelength of 320-400 nm (exact wavelength not specified in submission). Under white light, the beetle’s abdomen, pronotum, and head are a metallic dark green while its elytra are a metallic red/orange color. A crane fly (family *Tipulidae*) was found to fluoresce green on its head, thorax, and end of abdomen under an excitation wavelength of 400nm. Under white light, the insect appeared a beige color.

### Phylum Basidiomycota

A mushroom known as Coker’s amanita (*Amanita cokeri)* was found to fluorescence green on the remnants of its universal veil on the mushroom cap under an excitation of 400 nm. Under white light, these remnant pieces were white. A mushroom of the genus *Ganoderma* exhibited bright red fluorescence on the underside of its cap (which was a cream color in white lighting). There was little to no fluorescence on the top of the mushroom and a slight green, fluorescent ring on the margin of the mushroom (i.e. around the edge of the cap). Both the red and green fluorescent traits were observed in the *Ganoderma* mushroom when excited with light at 395 nm. A mushroom in the genus *Russula* was found to uniformly fluoresce green under a wavelength of 390 nm*. Under white light, this mushroom displayed a white stem and a light brown cap.

### Phylum Chordata

A pet ball python *(Python regius)* in a submitted observation emitted orange fluorescence, as seen especially on the lighter brown scales of its pattern. This fluorescence occurred when the observer placed the snake underneath a wavelength of 400 nm. In white light, the ball python had a combination of darker black and lighter brown scales forming its scale pattern. A green sunfish *(Lepomis cyanellus*) displayed potential fluorescence on some scales around its gills when excited with a light of 320-400 nm (exact wavelength not specified in submission). The exact emission color from this potential fluorescence occurrence is unclear. We can neither confirm nor deny fluorescence observations in *L. cyanellus* from the documented submission. A Mediterranean house gecko *(Hemidactylus turcicus)* fluoresced a neon green when exposed to light at 400-415 nm. The observer documented bright green fluorescence of its bone visible through skin, with areas on its skull and spine appearing the brightest. In natural light, the gecko appeared a pale brown color with dark markings scattered across its body. The Squirrel tree frog (*Hyla squirella*) had fluorescence displayed as a green stripe from snout to inner thigh while excited by light at 400-415 nm. In natural light the squirrel tree frog appeared a brown and green color with dark markings all over its body. The common Mexican treefrog *(Smilisca baudinii)* has green, fluorescent spots on its flanks and near its inguinal area under light at 400-415 nm. Under white light, the Mexican treefrog appears brown with darker patterned markings.

### Phylum Mollusca

A shell of the species of predatory snail named the shark eye (*Neverita duplicata*) was submitted to *Finding Fluorescence.* The shell was found to have a white/pink fluorescence in the spiral of its shell under a wavelength of 390 nm. Under white light, this spiral appeared white in color.

### Phylum Tracheophyta

A green bell pepper (*Capsicum annuum*) fluoresced orange and pink when held under an excitation wavelength of 390 nm. The pink fluorescence was especially concentrated toward the stem of the bell pepper, while the orange fluorescence was uniformly distributed across the body of the pepper. Under white light, the bell pepper and stem was dark green.

### Unidentified taxa

Observations from many different taxa were submitted to *Finding Fluorescence* and much effort was placed in concluding accurate identification for each organism. The subsequent section is designated for taxa that we were unable to narrow down to family classification, either because it would be impossible for such a narrow classification with the given photo submission or because our confidence in possible identifications was not strong. Identifications of these taxa represent our best estimates, given the guess from the submitter and the best judgement of our biologist collaborators. These submissions are grouped into categories: fungi, invertebrates, lichen, and plants. The observations are listed in chronological submission date order under each category. The full detailed submissions are available on the *Finding Fluorescence* website.

### Fungi

An observation of a mushroom submitted on 17 July 2020 from Richfield Township, Michigan, USA displays a green, fluorescent ring on the margin of the mushroom (i.e. around the edge of the cap) and in the center of the cap under wavelengths of 390 nm. Under white light, the mushroom is a solid white color with light brown speckles in the center of the cap. We could not conclude any confident taxa identifications for this submission.

### Invertebrates

A submission of a caterpillar documented on 26 May 2021 from Tallahassee, Florida, USA displays reported uniform light gray-green fluorescence under a wavelength of 365 nm. The submitter remarked that fluorescence was not observed under 395 nm. From the photo available, we are not able to determine if fluorescence is present. We do not have any confident identification proposals for the caterpillar. Another observation of what a submitter believed to be an aphid was documented on 8 July 2020 in Livonia, Michigan, USA. In this photo, an insect on a leaf is seen to fluorescence a uniform bright green under wavelengths of 395 nm. Under white light, the insect is solid white and is surrounded by material coated on the leaf that appears cotton-like. We are confident the submission is not an aphid. The best estimate for the insect described is a planthopper nymph (genus *Flatormenis*). A submission of a juvenile crab on 16 August 2020 from Dennis, New Jersey, USA reveals green fluorescence on the tips of the crab’s pereiopods (i.e. walking legs) under a wavelength of 390 nm. Under white light, the crab is a light white color. The suggested identification from the submitter is a juvenile spider crab, but we are confident this is not the correct identification. Our best assignment is a crab in the order Decapoda. Another submission of a millipede on 21 August 2020 from Tallahassee, Florida, USA displays bright green fluorescence across its entire surface under an excitation wavelength of 395 nm. The fluorescence is brightest on the underside of the millipede. No photos of the organism in white light were included. The millipede could possibly belong in the family *Xystodesmidae*, but we are not confident in this suggestion. A submission of a moth sent on 15 September 2020 from Groningen, Netherlands displays green fluorescence on the eyes of the insect under an excitation wavelength range of 320-400 nm (exact wavelength not specified in submission). Under white light, the moth is a dark brown, as are its eyes. A possible identification for the submission is *Noctua pronuba*; however, it is impossible to know for certain from the photo alone. Another submission of a caterpillar submitted on 20 September 2020 from Groningen, Netherlands displays bright green fluorescence on the entire surface of the insect. Under white light, the caterpillar is hairy and white. We are confident this caterpillar belongs in the order Lepidoptera, but no identification possibilities are available beyond that.

### Lichen

A submission of what appears to be lichen on a rock on 12 July 2020 from Fort Gratiot Township, Michigan, USA shows bright orange fluorescence under an excitation wavelength of 395 nm. Under white light, the possible lichen is light green in color. No suggested identifications are available for this submission. Another submission of what appears to be a lichen on 20 September 2020 from Groningen, Netherlands displays bright orange/yellow fluorescence when excited in the range of 320-400 nm. Under white light, what is possibly the rhizine (i.e. hair like attachment structures) is an orange color, while the possible cortex (i.e. top) of the possible lichen is a light grey color. This submission could possibly be a fruticose lichen, but we are uncertain in this suggestion.

### Plants

A submission of what appears to be roots of a plant on 8 July 2020 from Livonia, Michigan, USA fluoresce a bright green color under an excitation wavelength of 390 nm. Under white light, it is not able to be discerned what the suggested plant roots look like. No identifications are possible from the photo submission. A submission of a vascular plant leaf on 31 July 2020 from Campton Hills, Illinois, USA reveals slight red fluorescence under a wavelength range of 320-400 nm (exact wavelength not specified in submission). From the photo alone we are unable to provide a possible identification.

## Discussion

### Prior relevant documentations of biofluorescence

Many new observations of fluorescence were described through *Finding Fluorescence.* However, some of the submissions sent in were not the first documentation of fluorescence in that species. Here, we report on prior fluorescence documentation found in related taxa (up to same family) if applicable to the submissions (Table 3). An observation was considered novel if new fluorescent observations (such as location or color) were found in the organism, or if the excitation wavelength range used was different from past studies. Once again, the related taxa will be grouped by phylum.

### Phylum Arthropoda

A submission to *Finding Fluorescence* displayed distinct fluorescence colors of blue and yellow separately on cocoons of the domestic silk moth (*Bombyx mori*) under a wavelength of 395 nm. A past study had also documented distinct yellow-white and blue-purple fluorescence observations on the moth’s cocoon fibers under a different excitation wavelength of 360 nm [22]. Another study documented the separate blue and yellow fluorescence in the cocoon fibers under a slightly different excitation of 365 nm [23]. We consider the submission sent in to *Finding Fluorescence* a new observation, as the excitation wavelength range is different from previous findings. The submission we received to *Finding Fluorescence* also showed the abdomen of the *B. mori* to fluoresce yellow under an excitation wavelength of 395 nm. No studies examining the fluorescence of the abdomen of the moth itself could be found. Another submission of the blue dasher dragonfly (*Pachydiplax longipennis*) revealed no fluorescence under an excitation wavelength of 395 nm. However, blue fluorescence has been recorded in *P. longipennis* along its thorax and wing hinges under excitation of 310-340 nm [24]. This suggests the fluorophore producing fluorescence in this species likely has a peak emission of 310-340 nm or lower, and future studies should test fluorescence in *P. longipennis* with even shorter wavelengths. A submission of the Oslar’s eacles moth (*Eacles oslari*) displayed yellow fluorescence on the abdomen of the moth under a wavelength of 395 nm. Prior orange and brown-lavender fluorescence in an unspecified location has been documented before in a related moth *Eacles imperialis* under a wavelength range of 320-400nm [25]. The submission we present for *E. oslari* appears to be a new fluorescence documentation. No proposed function of this fluorescence occurrence could be found.

### Phylum Basidiomycota

A submission to *Finding Fluorescence* of a mushroom in the genus *Russula* revealed green fluorescence on the entire surface of the mushroom under an excitation wavelength of 390 nm. This is not the first fluorescence documentation for this genus. Many species in the genus *Russula* have been found to fluoresce under an excitation wavelength range of 320-400 nm (exact excitation wavelength, emission color, and location not specified) [26]. It is known that fluorescence in various *Russula* species is due to specific water-soluble pteridines called russupteridines [33]. Violet-blue fluorescence is due to pro-lumazine, and yellow fluorescence occurrences in *Russula* are due to russupteridine-yellow [33].

### Phylum Chordata

A submission to *Finding Fluorescence* of a Mediterranean house gecko *(Hemidactylus turcicus)* showed green fluorescence of the gecko’s skull and spine through its skin when exposed to light at 400-415 nm. The submission to *Finding Fluorescence* appears to be the first fluorescence documentation for the species *H. turcicus* in the wavelength range of 400-415 nm. There were previous fluorescence documentations in the same genus *Hemidactylus*. Green fluorescence of the skull, spine, and appendages of a closely related species *Hemidactylus frenatus* has been documented before under a different excitation wavelength range of 395-400 nm [27]. Under a wavelength of 395 nm, it was found that another related species *Hemidactylus platyurus* fluoresced blue along the lower jawbone [28]. Another related gecko (*Pachydactylus rangei)* within the same family *Gekkonidae* fluoresced neon green along the skin around the eyes and along the flanks under wavelength 365 nm [29]. It has been discovered that bones in humans naturally fluoresce due to type I collagen, which is the most abundant collagen in connective tissues such as bone [34]. Assumably, gecko bones would also contain type I collagen, which may explain the physical reason why the fluorescence is observed in the bone of the gecko. A submission to *Finding Fluorescence* of the common Mexican treefrog *(Smilisca baudinii)* shows green, fluorescent spots on its flanks and near its inguinal area under light at 400-415 nm. Green fluorescence along the arms and inguinal area of *S. baudinii* has been documented prior under the same wavelength of 400-415 nm [30].

### Phylum Tracheophyta

A submission to *Finding Fluorescence* of a green bell pepper (*Capsicum annuum*) fluoresced orange and pink when held under an excitation wavelength of 390 nm. Fluorescence has been recorded before for *C. annuum.* In prior studies, the pepper fluoresced blue under excitation wavelengths of 365 nm [31]. Blue fluorescence, low-green, and chlorophyll fluorescence were also documented in *C. annuum* at a maximum excitation wavelength of 440 nm [32].

### Potential ecological relevance of fluorescence

The ecological significance that biofluorescence holds for an organism is an area of ongoing study across many taxa. Many studies in fluorescence have been concentrated on marine organisms, but fluorescence observations in terrestrial organisms is an area of relatively untapped potential. While we sought to document new findings of fluorescence across various organisms, we also considered potential roles for the fluorescence present across the many taxa presented in this paper. As in previous sections, commentary on fluorescence in some of the organisms documented will be formatted according to phylum groupings. It should also be considered that not everything that fluoresces may possess an ecological function of that fluorescence – it may simply be a byproduct of certain structures within that organism that possess such qualities (i.e. chemical structures known as fluorophores), but it may not serve the organism in an additional way. Where no fluorescence commentary is provided for a particular organism present in our observations, it is because there is insufficient literature on how coloration plays a role in visual communication. We suggest that these unmentioned fluorescent organisms be considered for future work in visual aspects of their communication and explored further.

### Phylum Arthropoda

Fluorescence in insects was first documented in the 1920s [35]. Since then, insect fluorescence has been observed across many organisms, but much more needs to be studied in this group. We have gained fluorescence submissions from many insects such as beetles, moths (caterpillars and adults), dragonflies, millipedes, crane flies, and spittle bugs.

Interestingly, the fluorescence emissions of the cocoon fibers of the domestic silk moth (*Bombyx mori*) are determined by the sex of the moth inside each cocoon. [22, 23]. Each of the fluorescent cocoons submitted to *Finding Fluorescence* were distinct in their emissions (e.g. one cocoon is blue, and another cocoon is yellow). It has been found that the fluorescence emission colors are sex-related, and that blue fluorescence indicates a female silkworm and yellow fluorescence indicates a male silkworm [23]. The difference in fluorescence emissions is largely due to distinct transportation patterns and accumulation of fluorescent pigments in the midgut and in the silk glands for the separate sexes [23]. Whether the fluorescence of the cocoon fibers has any ecological function or is simply due to fluorophore presence in the midgut is unknown currently. Moths (and other insects such as bees and butterflies) possess the ability to see ultraviolet light [36], and future studies should aim to address such questions of if there is any purpose of having fluorescence in cocoon fibers of moths given their absence of parental care.

We have also gained fluorescent submissions from arthropod crustaceans such as mole crabs and true crabs. In the case of the mole crab *E. talpoida*, we propose that the fluorescence found along its legs and the crevices of its exoskeleton are a biproduct of fluorophores present in the structural proteins of such areas. In the case of the unidentified submission of a true crab we received, fluorescence was only observed at the tips of its pereiopods. We also suspect this to be due to fluorophores present at that area.

Due to the lack of information on fluorescence in insects, the exact ecological roles and abundance of fluorescence across insect groups cannot be generalized [37]. Further studies need to be conducted before anything more can be said about the fluorescence of other arthropod submissions to *Finding Fluorescence.* We suggest studies examining the fluorescence of Coleoptera, Lepidoptera, Hymenoptera, Odonata, and Decapoda would be good places to start, as these orders lack robust fluorescence documentations.

### Phylum Basidiomycota

We have fluorescent observations from three distinct genera of mushroom (*Amanita, Ganoderma,* and *Russula*). The pattern of fluorescence differs in each genus received. In the case of the *Amanita* mushroom, fluorescence was only present on the universal veil remnant; in the *Ganoderma*, the margin of the mushroom and the underside both emitted different fluorescent colors; and the *Russula* displayed a uniform emission. Insects are one of the many taxa which possess the ability to see into the UV spectrum [38], and it has been proposed that perhaps fluorescence serves to attract such invertebrates for aid in spore dispersal [39]. Future studies should aim to further test this proposal and investigate possible ecological relevance in fluorescence patterns and colorations in Basidiomycota.

### Phylum Chordata

*Finding Fluorescence* received fluorescence observations for a ball python, Mediterranean house gecko, squirrel tree frog, and the Mexican tree frog. A study examining the fluorescence function in pitvipers found an association between fluorescence presence, caudal luring (i.e. moving one’s tail in a manner to attract prey), and conspicuous tail coloration [6]. Further studies should be conducted to examine if the presence of tail-tip fluorescence is seen in other snake families which exhibit caudal luring. Considering the potential fluorescence relevance in the case of the *Finding Fluorescence* submission of *Python regius*, it should be noted that the snake is a ground-dwelling species with no caudal luring behavior and with different overall ecology than pitvipers. A suggested potential relevance for the orange fluorescence documented in *P. regius* could be to aid in its camouflage. The orange fluorescence could more closely match the background of red fluorescence of leaves in the forest, as chlorophyll in plants is found to fluoresce red under excitation wavelengths of 400-700 nm [40]. Further studies should aim to test such hypotheses of other ground-dwelling snakes to test for possible camouflage aid.

It has been suggested that fluorescence in geckos can aid in intraspecific signaling or identification by accentuating certain aspects of the gecko, such as body markings or its skeletal elements [41, 42]. In our *Finding Fluorescence* observation of the Mediterranean house gecko (*Hemidactylus turcicus*), green fluorescence of the skull is observed under 400-415 nm of excitation wavelength. In previous findings, fluorescence has been observed in at least two other members of the same genus: *H. frenatus* and *H. platyurus*. In the case of *H. frenatus,* blue fluorescence was observed on the skull, spine, and appendages. For *H. platyurus*, green fluorescence was observed along the lower jawbone. Another gecko in the same family as *Hemidactylus,* the Namib sand gecko (*Pachydactylus rangei*) displays green fluorescence along the skin around its eyes and flanks. All three mentioned related species fluorescence in different locations under differing excitation ranges (Table 3). The different locations of fluorescence as well as the different excitation wavelengths observed in the different species could further support the idea of fluorescence playing a role in intraspecific communication. Geckos have been found to have trichromatic color vision and are especially sensitive to wavelengths in the green, blue, violet, and UV range [43–49]. The documented fluorescent differences in geckos may indeed serve as a way for species to recognize other members of their own species.

In the two frog submissions we received, it was shown that the squirrel tree frog displayed a green, fluorescent line from its snout to its inner thigh, while the Mexican tree frog displayed green fluorescence spots along the flank of its body and near its inguinal area (i.e. near the groin). It has been shown in squirrel tree frogs that females have a strong preference for presence of a relatively large lateral body stripe in males [50]. This lateral body stripe was what was shown to fluorescence in the observation submitted to *Finding Fluorescence.* Due to the role the stripe plays in sexual selection, the fluorescence presence of the stripe could be indicative of intraspecific signaling in squirrel tree frogs and be utilized by females during mate choice. In the context of the Mexican tree frog submission we received to *Finding Fluorescence*, we propose its fluorescence pattern near the inguinal area to play a role in intraspecific communication. Over half of the studies examining ecological function in anuran biofluorescence have suggested fluorescence as a mechanism to deter predation. However, a recent study experimentally testing this hypothesis has found no such effect of biofluorescence as an antipredator signal in Cope’s gray treefrog (*Hyla chrysoscelis*) [51]. This study is one of the first examining the relationship between fluorescence and predation. Across anuran families, evidence has been found that fluorescence presence might be a signal for intraspecific communication [52]. Evidence for ecological tuning (i.e. the specific response of a signal to its environment) has been supported when examining fluorescence over 13 anuran families [52]. Future studies should aim to further explore the roles of alternate functions of fluorescence in frogs as well as the potential role in predator deterrence in other organisms.

### Phylum Mollusca

We received one submission in the phylum Mollusca in which a white/light-pink fluorescence was observed in the spiral of the shell of a predatory snail called the shark eye (*Neverita duplicata*). An extensive study examining the fluorescence in mollusk shell pigments suggests differing compounds allow for fluorescence in the patterns of mollusk shells [53]. The fluorescent pigments uroporphyrin I and III were identified to fluorescence red under UV light in some species of marine gastropods [53]. It is also suggested that fluorescence is only due to compounds located in the top layer of the shell and that metal ions may affect coloration of the pigment in the shell [53]. However, the fluorescence in the submission sent to *Finding Fluorescence* seems to be stemming from the spiral of the shell, not any type of coloration pattern. The exact nature and possible function (if any) of this fluorescence cannot concretely be speculated on at this time. The specific location of fluorescence could point toward a high concentration of different compounds or fluorophores in this area. Future studies should look at examining the specific compounds in different locations in molluscan shells and identifying fluorophores localized in those regions.

### Phylum Tracheophyta

We received a submission of fluorescence documented in a green bell pepper (*Capsicum annuum*). Measuring fluorescence in bell peppers has been proposed as a good method for examining pepper quality [31]. More specifically, it has been suggested that surface fluorescence in the pericarp as well as the cuticle of a pepper can be used to indicate postharvest quality [54]. As an example of how fluorescence can potentially indicate traits in peppers, it was found that the peppers which fluoresced darker were likely to have thicker cuticle and epidermis with a more uniform outer wall thickness [54]. The fluorescence in the bell pepper may simple be due to fluorophores present within the plant’s tissue, or perhaps the fluorescence may serve to attract insects or animals for seed dispersal purposes. Prior studies have looked at fluorescence in bell peppers from an agricultural perspective, but future studies should examine how this fluorescence in peppers may interact with the natural world.

### Significance of newly discovered fluorescence via community science

Our research began by documenting and assessing new occurrences of biofluorescence uploaded as community science data to the *Finding Fluorescence* resource. We verified and compared new biofluorescent accounts to the literature to provide insight into the role of biofluorescence across taxa. *Finding Fluorescence* is the first community science database to collect observations of biofluorescence in organisms. There are, of course, other collective studies examining fluorescence across organisms. For example, studies have sought to document larger fluorescence observations in amphibians [55], fish [1], Catsharks [2], and rodents [56]. These reviews differ in ours in that 1) they are not community science submissions, 2) they are more specific in study taxa, and 3) they are at different scales than our study. The study examining fluorescence in fish looked at 180 species across 16 orders [1]. The survey examining fluorescence in amphibians and salamanders examined eight families of salamanders, five families of frogs, and one family of caecilians [55]. To reconstruct a phylogeny of biofluorescence of catsharks, 77 taxa (including 74 Chondrichthyes representing 51 families) were analyzed [2]. Museum specimens of rodents from 28 genera (order Rodentia) were examined for fluorescence in the rodent study [56]. In the case of our study, we took a broad approach and analyzed and confirmed novel fluorescence submissions across five different phyla, 15 families, and 15 species. While our study is not as in-depth within any specific taxa, we believe presenting new documentations via community science involvement is an exciting new avenue to engage the broader public in science involvement and intrigue. Our study is also unique in that many of the novel observations of fluorescence presented have differed in excitation wavelength, emission wavelength, and locations than previous findings of the same organism. Understanding and documenting fluorescence at different levels of excitation is an untapped area of research that we hope to further with the continuation of the *Finding Fluorescence* community science database.

### Improvements to resource

We identified several areas of *Finding Fluorescence* that have since been improved based on our experience with assessing community science submissions. We have now designated all submission fields to be required (i.e. the observer cannot leave any fields blank when submitting an observation). By having such fields required, it helps in ensuring a higher quality of submission data to work with. Previously, the section for the community scientist to submit their proposed common or scientific name for the organism was an optional field. In some observations where the photo was blurry or taken from far away, the proposed identification of the organism was left blank, and we were unable to propose any type of identification with the given information. It would have been beneficial to have the suggestive identification from the submitter be required to have aided in organism identification. Going forward, we hope this required field will minimize submissions not being able to be verified and identified.

As well, we have now added clarifying methodology for photo submissions, including specific instructions that emphasize the need for photos in natural/white light and a request that all images capture the entire organism (as possible) and to capture photos of smaller organisms (i.e. small arthropods, small plants) at a max distance of around six inches away from the organism. These added instructions will aid in visual organism identification and allow for more possible organism identifications. Additionally, another section for photo submission that requests multiple angles of the organism to assist in identifying organisms that may have recognizable markings on specific regions of their surfaces has been added. A section which contains contact information for the submitter has since been required to verify information (if needed). Under this field, we have also included a question asking the submitter they to being recognized for their fluorescence discovery. We hope that by offering the opportunity for acknowledgement in scientific discovery will increase community investment in science early on as well as aid in documenting even more novel observations. This field was present before, but optional, and we have only received one submission since the start of the site that contains personal information of the submitter. We have now made this specific field viewable only to the managers of the *Finding Fluorescence* site to protect personal information of submitters online. We have also included an instructional photo of an example of how to take a detailed and quality photo of fluorescence directly on the page the submitters upload observations to. Previously, the photo could be found elsewhere on the site, but we hope that by including the photo again on the submission page directly, the submitter will have a better idea of the criteria which should be met with their submission.

## Acknowledgements

We gratefully acknowledge all the community science members who submitted their fluorescence observations to *Finding Fluorescence*, including Amanda Sakimura who sent in museum specimens of *Eacles oslari* and *Bombyx mori*. We thank Brian Inouye and Nora Underwood for their aid in identifying our presented Basidiomycota submissions (*Ganoderma* mushroom*, Russula* mushroom, and *Amanita cokeri*) and the *Tipulidae* family crane fly. We are also grateful for the FSU Ecology and Evolution Reading Discussion Group (EERDG) for help in creating and funding *Finding Fluorescence* education and outreach materials.

## Author contributions

**Hannah Burke:** investigation (lead); methodology (supporting); visualization (lead); validation; writing – original draft preparation (lead); writing – review and editing (equal). **Lauren Serrano:** investigation (supporting), writing – original draft preparation (supporting). **Emily Lemmon:** project administration (supporting); supervision (supporting); writing – review and editing (supporting). **Courtney Whitcher:** conceptualization; methodology (lead); project administration (lead); resources; supervision (lead); writing – review and editing (equal).

